# Reliable prediction of protein-protein binding affinity changes upon mutations with Pythia-PPI

**DOI:** 10.1101/2024.10.28.620752

**Authors:** Fangting Tao, Jinyuan Sun, Bian Wu, George F Gao

## Abstract

Protein-protein interactions are essential for numerous biological functions, and predicting binding affinity changes caused by mutations is crucial for understanding the impact of genetic variations and advancing protein engineering. Although machine learning-based methods show promise in improving the prediction accuracy, the scarcity of experimental data remains a significant bottleneck. Here, we utilized multi-task learning and self-distillation to overcome the data limitation and improved the accuracy of deep learning-based protein binding affinity prediction. By incorporating a mutation stability prediction task, the model achieved state-of-the-art accuracy on the SKEMPI dataset and was subsequently used to predict binding affinity changes for millions of mutations, generating an expanded dataset for self-distillation. Compared with prevalent methods, Pythia-PPI increased the Pearson correlation between predictions and experimental data from 0.6447 to 0.7850 on the SKEMPI dataset, and from 0.3654 to 0.6051 on the virial-receptor dataset. These findings demonstrated Pythia-PPI to be a valuable tool for analyzing the fitness landscape of protein-protein interactions. We provided a web server at https://pythiappi.wulab.xyz for easy access to Pythia-PPI.

## Introduction

Proteins, as essential macromolecules, play fundamental roles in many physiological functions within organisms [1]. Biological processes such as signal transduction, immune response and viral adhesion rely on the protein-protein interactions (PPIs) [2, 3, 4]. Therefore it is of fundamental interests to understand how mutations perturbate the thermodynamic properties of PPIs to deepen our insights into pathogenic genetic variations or optimize therapeutic proteins [5, 6]. The strength of PPIs is typically evaluated through binding affinity, commonly defined by the Gibbs free energy (ΔG) [7]. Assessing the effect of mutations can be achieved by measuring the difference in ΔG between the mutant and wild-type, referred to as ΔΔG, which can evaluate the effect of the mutation on PPI binding affinity or protein stability. Although experimental methods including isothermal titration calorimetry [8] and the yeast two-hybrid system [9] can be used to measure ΔΔG values in PPIs, these methods are generally time-consuming and labor-intensive. Given these limitations, a reliable way to predict the ΔΔG of protein binding due to mutations is urgently needed.

Computational methods can be classified into two categories: physics-based methods and machine learning-based methods. The former, such as Rosetta [10] and FoldX [11], utilize energy functions, which consider physical interactions such as hydrogen bonds and van der Waals interactions between atoms. Although these methods can provide interpretable results, they require extensive sampling of conformation space and suffer from limited accuracy of energy functions due to approximate assumptions in dealing with anisotropic interactions [12].

In contrast to physics-based methods, machine learning-based methods are mainly data-driven. Although many machine learning algorithms, including artificial neural networks, used tools from physics, neural networks capture data representations through various layers of abstraction and identifies complex patterns within data rather than relying on human-defined rules [13, 14]. Deep learning has assisted scientific discoveries in the field of protein science, such as accurate structure prediction with AlphaFold2 [15] and advanced the development of computational protein design [16]. There are also attempts to address the binding affinity prediction challenge using deep learning. Graph neural networks such as DDGPred [17] require Rosetta modeled mutant structures and energy terms. Convolutional neural network exemplified by the Binding Oracle [18] required voxelization of protein local structure. Beyond these computational requirements, a fundamental challenge in the prediction of ΔΔG is the scarcity of labeled data. This scarcity makes it difficult to fully capture the complex structure-function relationships within protein-protein complexes, which results in a low success rate for optimizing antibodies [17]. Overcoming data limitations and developing reliable, efficient deep learning-based predictors for ΔΔG values in PPIs is essential for further applications.

In this study, we developed Pythia-PPI and achieved significantly higher prediction accuracy of protein-protein binding affinity changes upon mutations. Firstly, a vanilla Pythia-PPI from the pre-trained Pythia model using multi-task learning and enabled simultaneous prediction of mutation impacts on both protein stability and protein-protein binding affinity. By leveraging the efficiency and precision of this baseline model, we generated mutation predictions for all protein interaction interfaces within the SKEMPI dataset [23], expanding the training set for PPI-related mutations to near 400 thousand. With this augmented dataset and multi-task learning, Pythia-PPI achieved a Pearson correlation of 0.785 on the SKEMPI dataset. Additionally, Pythia-PPI provides exceptional throughput, capable of processing over 10,000 mutation predictions per minute, facilitating large-scale point mutation analyses across the human interactome. For broader accessibility, predictions for protein complexes can be freely conducted on our web server at https://pythiappi.wulab.xyz, and the source code is openly available at https://github.com/Wublab/pythia_ppi.

## Results

### Pythia-PPI architecture

Pythia-PPI consists of two modules: a pre-trained structure graph encoder module and a ΔΔG prediction module. Employing a k-nearest neighbor (k-NN) graph, Pythia-PPI transforms the local structure of a protein or protein-protein complex into a graph representation (Figure 1A). Each amino acid acts as a node, establishing connections with its 32 closest amino acids based on the Euclidean distance of the C-alpha atom. The input of the structure graph encoder incorporates the type of each amino acid through one-hot encoding, while representing backbone dihedral angles (φ, Ψ, and ω) using sine and cosine functions as node features. Regarding edge features, we consider distances between the five backbone atoms: C-alpha, C, N, O, and C-beta, along with sequence positions and chain information. Node and edge input features are converted to embeddings through the structure graph encoder, identical to the pre-trained Pythia [24]. This encoder includes three attention message passing layers (AMPLs) that generate hidden embeddings and a linear layer that generates probabilities for the 20 amino acids (Figure 1B).

**Figure 1.**
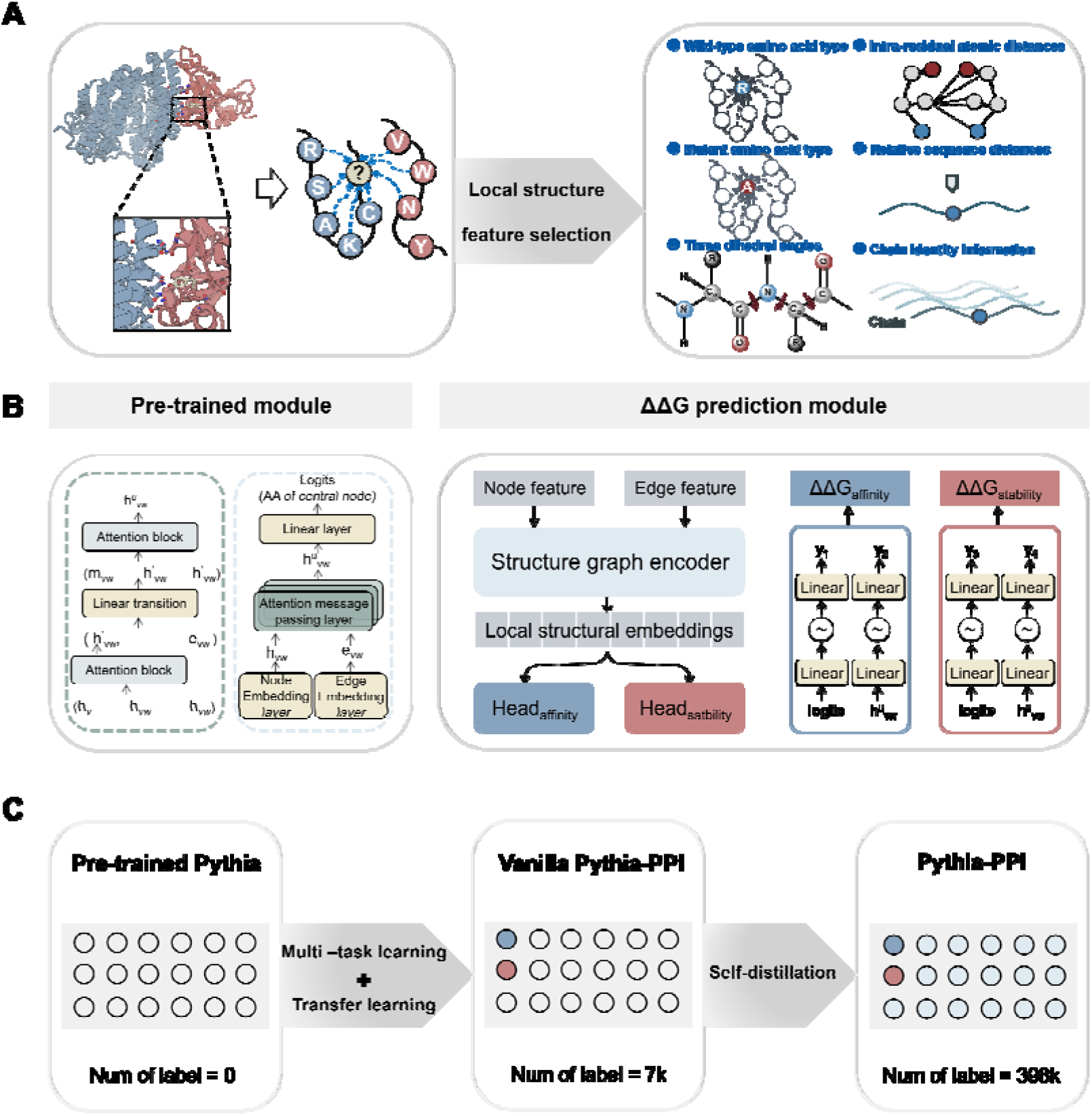
Overview of Pythia-PPI. (A) Pythia-PPI receives a local structure of either a protein-protein complex or a protein as input, depicted as a k-NN graph derived from the Euclidean distances of C-alpha atoms. (B) The architecture of Pythia-PPI, trained on representations extracted from a pre-trained model, aimed at predicting binding affinity changes of PPIs or stability changes of proteins caused by single-point mutations. (C) The process of developing the Pythia-PPI with multi-task learning and data augmentation.

In Pythia-PPI, hidden embeddings are extracted from each of the AMPLs, along with amino acid probabilities, focusing on features of residues that are either wild-type or mutant. Utilizing the message-passing mechanism inherent in Pythia, information about neighboring residues of both wild-type and mutant residues is captured within a structural context. These embeddings are then combined with corresponding probabilities from the pre-trained module to form the input vector for the ΔΔG prediction module. Pythia-PPI employs a combination of transfer learning and multi-task learning, sharing the structure encoder layer across two tasks: predicting changes in PPI binding affinity and protein stability upon mutations. The ΔΔG prediction module comprises two heads, named head_affinity_ and head_stability_, whose selection is determined by the data source of input features. The chosen input vector determines its allocation to either head_affinity_ or head_stability_. Each head consists of two compact multi-layer perceptrons (MLPs), aimed at producing ΔΔG values.

As shown in Figure 1C, we first evaluated the zero-shot transferred prediction of self-supervised models to find a model learned a potentially better representation with balanced computation costs. Then, we improved the accuracy on protein binding affinity prediction task by multi-task supervised fine-tuning. With an accurate predictor developed, we attempted to improve the model with self-distillation strategy.

### Supervised fine-tuning largely improved prediction accuracy than zero-shot transferring for binding affinity prediction

Previous studies have shown that protein sequence likelihood models can be effectively generalized to the prediction the impact of mutations on thermal stability in a zero-shot manner.

Our previous study established a physical relationship between amino acid probability distributions within a protein’s structural context and the thermostability changes associated with protein mutations. Building on this finding, we developed the Pythia model, which achieved state-of-the-art performance in unsupervised thermostability prediction. In this study, we first evaluated Pythia’s zero-shot predictive capacity for ΔΔG changes in protein-protein interactions, utilizing the SKEMPI dataset [23] as a benchmark. The Pearson correlation of Pythia is 0.3782, out-performed the MIF model [25], which is also a self-supervised structure-aware sequence likelihood model (Table S1). In addition, as depicted in Figure 2A, Pythia exhibits a significantly faster computational speed compared to MIF, which will be beneficial to apply Pythia-based methods from large scale analysis.

**Figure 2.**
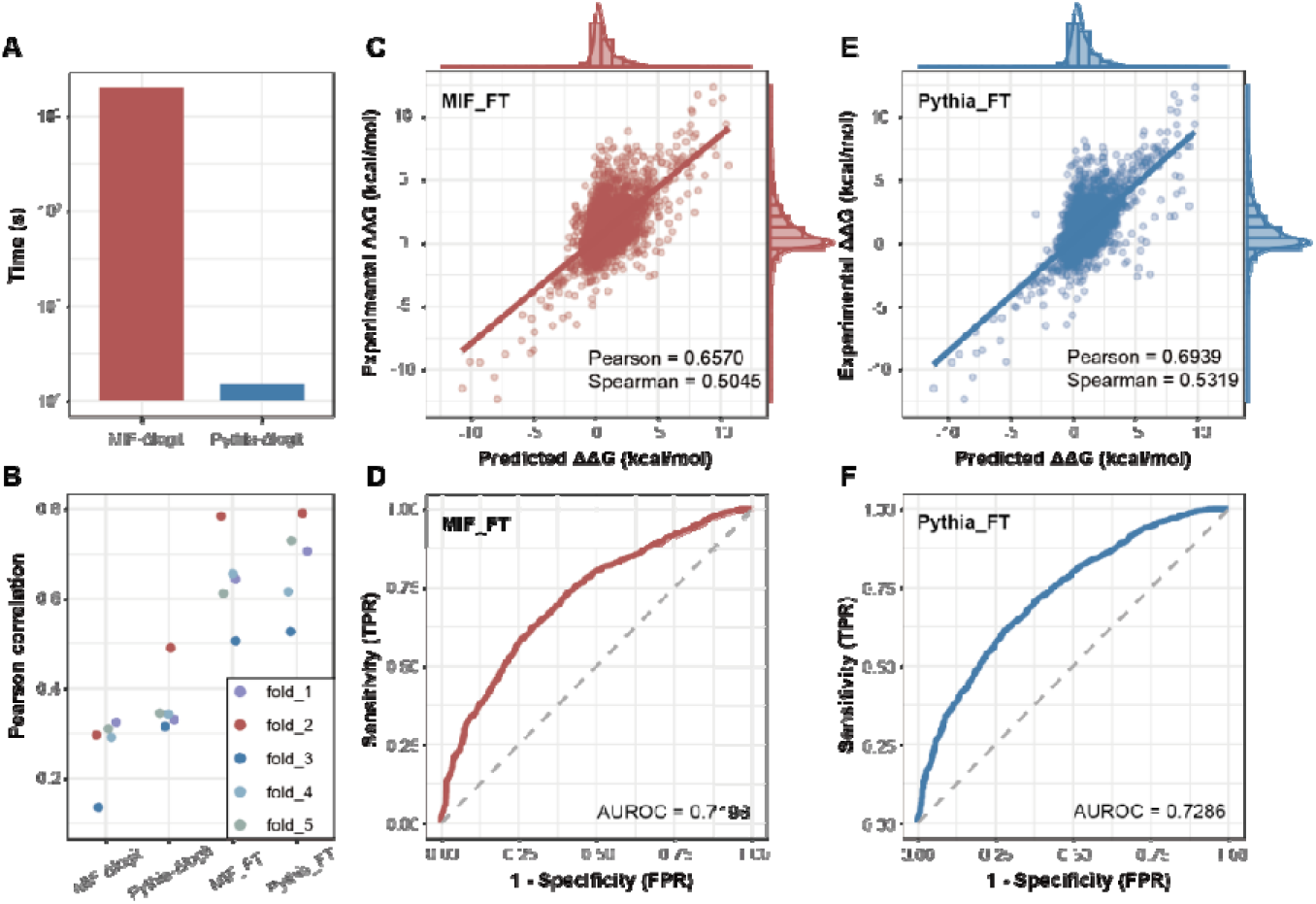
Performance of pre-trained and fine-tuned Pythia and MIF. (A) Comparison of the computing speed between two methods using all possible single-point mutations based on the PDB structure 6M0J. (B) Pearson correlation calculated on predictions for models trained on the SKEMPI dataset by five-fold cross-validation. (C–F) Scatter plots and ROC curves of MIF_FT and Pythia_FT predictions versus experimental values on the SKEMPI dataset, where (C) and (D) show MIF_FT results, and (E) and (F) show Pythia_FT results.

Both Pythia and MIF were fine-tuned on the SKEMPI dataset, resulting in the fine-tuned models Pythia_FT and MIF_FT. After fine-tuning, both models demonstrated improved prediction accuracy for protein binding affinity in five-fold cross-validation compared to their zero-shot predictions (Figure 2B). When comparing the performance of the two, Pythia_FT consistently outperformed MIF_FT on the SKEMPI dataset: MIF_FT achieved Pearson and Spearman correlations of 0.6570 and 0.5045, with an AUROC of 0.7198 (Figures 2C and 2D). In contrast, Pythia_FT showed enhancements with Pearson and Spearman correlations of 0.6939 and 0.5319, and an AUROC of 0.7286 (Figures 2E and 2F). Given Pythia’s superior speed, zero-shot transfer performance, and fine-tuned prediction accuracy, we selected it as the basis for further multi-task predictive model development.

### Multi-task learning enhanced the accuracy of structure-wise binding affinity prediction

The prediction of stability changes upon mutations was introduced as another training task, to enhance the understanding of the relationship between structural and energy changes during the supervised learning. This could let the model to learn shared representations of common features between protein structure and thermodynamic parameters from both protein folding stability and protein-protein binding affinity. The FireProt DB contains experimental thermostability data, derived from published datasets and data extracted from recent literature manually [26]. This comprehensive database was carefully filtered to create the F3436 dataset suitable for machine learning, which contained 3,436 single-point mutations across 100 proteins [20]. By integrating the F3436 dataset, increases the representation of specific mutation types than using the SKEMPI dataset alone (Figure S1). For training, we combined data from four folds of the SKEMPI dataset with the F3436 dataset, holding out the remaining SKEMPI fold for validation.

Through grid search of the weights of stability and affinity losses, we identified an optimal stability-to-affinity loss ratio of 0.8:0.2, which maximized Pearson correlation (Figure S2). After incorporating the stability prediction task, the structure-wise prediction correlations improved noticeably, with Pearson correlation increasing from 0.4462 to 0.4778 and Spearman correlation from 0.4119 to 0.4486 (Figure 3A, Table S1). In five-fold cross-validation on the SKEMPI dataset, the vanilla Pythia-PPI achieved a AUROC of 0.7341 (Figure 3B), a Pearson correlation of 0.7092, and Spearman correlation of 0.5363 (Figure 3C). A strong Pearson correlation of -0.95 between the predicted ΔΔG of direct and reversed mutations demonstrated that the prediction of our model is almost anti-symmetric (Figure 3D). This indicates that our model has a lower risk of overfitting and improved generalization capability. Comparing with other state-of-the-art methods, the vanilla Pythia-PPI improved the per-structure Pearson correlation from 0.4448 to 0.4778, and the per-structure Spearman correlation from 0.4010 to 0.4486 (Figure 3E). This improvement may be due to the increased structural diversity in the training data introduced by the stability prediction task, enabling the model to better capture the relationship between protein structure and thermodynamic parameters.

**Figure 3.**
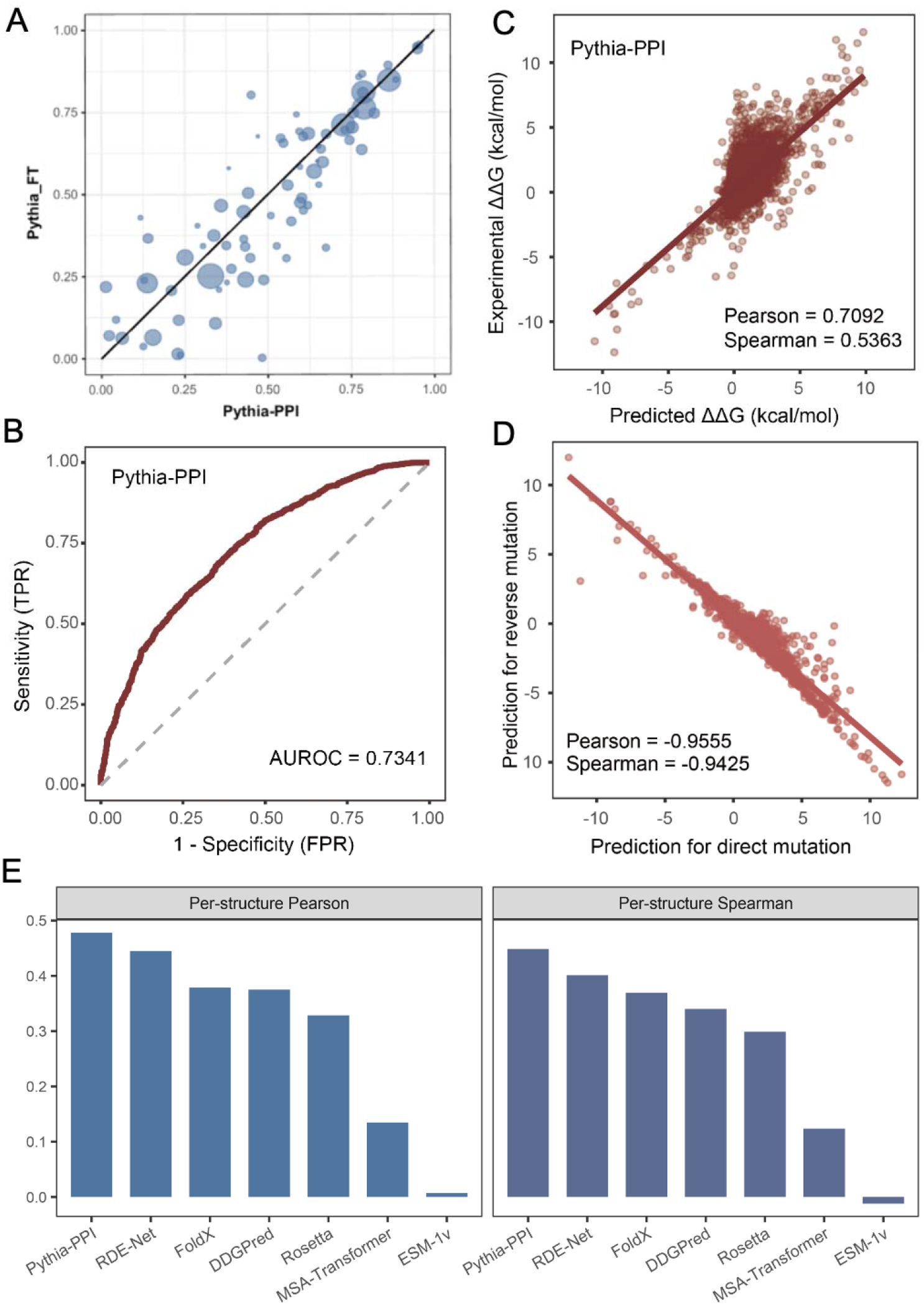
Performance analysis of the vanilla Pythia-PPI on SKEMPI dataset. (A) Comparison of per-structure Pearson values of Pythia_FT and the vanilla Pythia-PPI. (B) Receiver operating characteristic curve of the vanilla Pythia-PPI. (C) Scatter plot of experimental and predicted ΔΔG on the SKEMPI dataset. (D) Comparison of predictions for direct and reverse mutations. (E) Bar plot of per-structure Pearson and Spearman correlation on SKEMPI dataset of tested methods.

### Data augmentation boosted the prediction accuracy of binding affinity

To further expand the training data and enhance model prediction accuracy, we adopted a self-distillation strategy. Self-distillation involves training a model using its own predictions as additional labeled data, which can improve generalization by exposing the model to a broader distribution of examples. Using vanilla Pythia-PPI, we predicted the ΔΔG of saturated mutations across all amino acids at the interfaces of protein complexes in the SKEMPI dataset. This increased the training dataset for protein affinity prediction from around 4,000 to nearly 400 thousand samples. When projected into two dimensions using t-SNE on local structure embeddings from Pythia, the self-distilled samples exhibit broader and denser spatial coverage compared to the original SKEMPI dataset samples (Figure 4A). While most complex structures in the original SKEMPI dataset contain only dozens of labeled mutations, the self-distilled dataset includes between 200 and 2,000 mutations per complex (Figure 4B). With this augmented training data, we trained the final Pythia-PPI with the multi-task strategy as previous mentioned.

**Figure 4.**
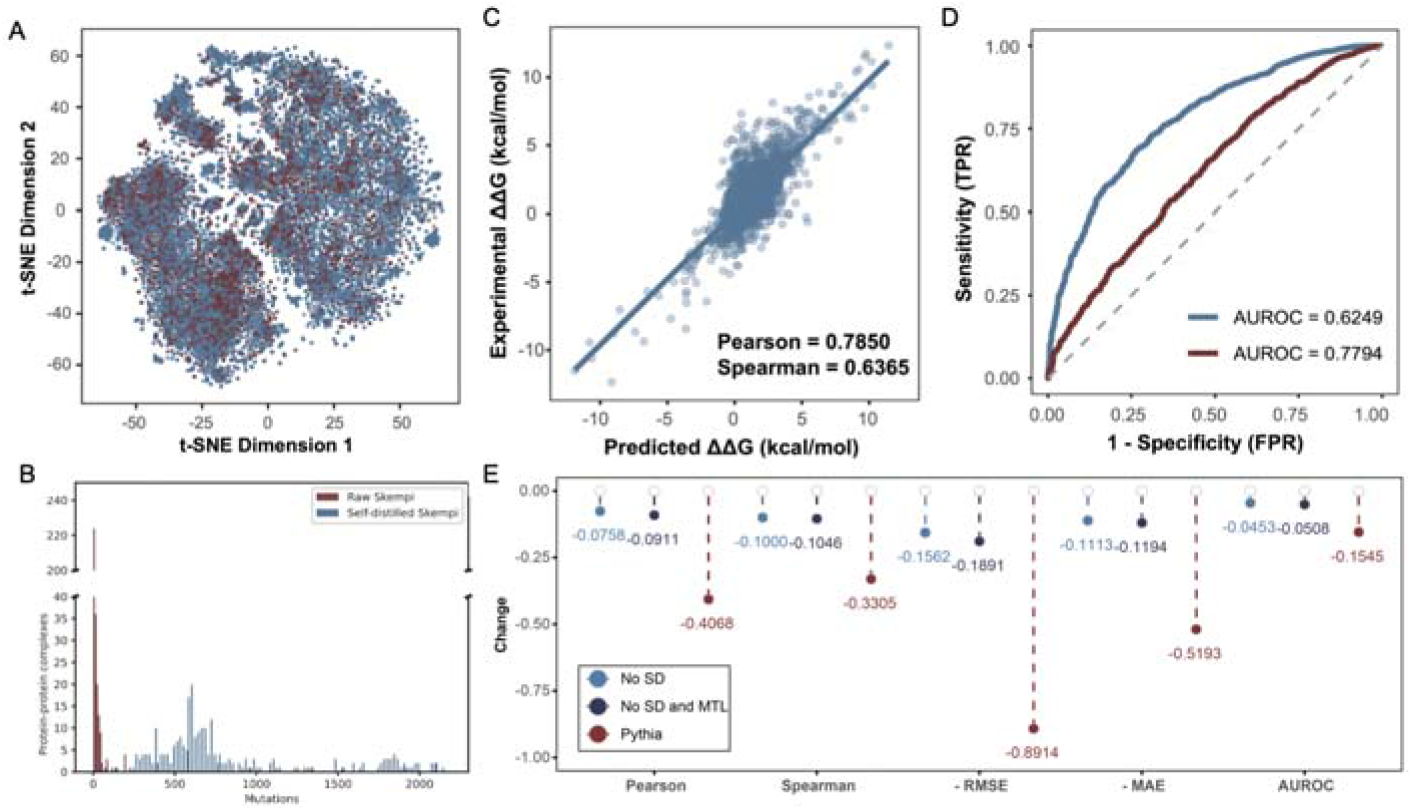
Performance analysis of Pythia-PPI based on the SKEMPI dataset. (A) 2-dimensional visualization of structural distribution of training data in SKEMPI experimental dataset (red dots) and SKEMPI augmented dataset (blue dots). (B) Bar plot of counts of complex structure versus mutation numbers. (C) Scatter plot of Pythia-PPI predictions versus experimental values. (D) Receiver operating characteristic curve of the final Pythia-PPI and the zero-shot transferred Pythia. (E) Reduced level of different ablation settings of metrics on the SKEMPI dataset.

In benchmark evaluations on the SKEMPI dataset, Pythia-PPI demonstrated robust predictive performance. As shown in Figure 4C, the model exhibited a Pearson correlation of 0.7850 between predicted and experimentally measured ΔΔG values, while achieving a Spearman correlation of 0.6365 that confirms its ability to accurately capture the relative trends in protein-protein binding affinity. Compared to the baseline model, Pythia-PPI achieved substantial improvement in discriminating stable versus unstable interactions, while Pythia achieved an AUROC of 0.6249 in zero-shot prediction, our specialized model reached an AUROC of 0.7794 (Figure 4D).

When compared with other state-of-the-art predictors, Pythia-PPI demonstrated superior performance across all key metrics (Table 1). Traditional physics-based approaches, exemplified by Rosetta and FoldX, achieved Pearson correlations of 0.31, while protein language models like ESM-1v [28] and MSA Transformer [29] showed limited predictive power with Pearson correlations values below 0.2. Pythia-PPI outperformed the previous state-of-the-art predictor RDE-Net [30] by improving the overall Pearson correlation from 0.645 to 0.785. The advancement was also evident in per-structure evaluation, where Pythia-PPI achieved a Pearson correlation of 0.567 compared to RDE-Net’s 0.445, suggesting enhanced generalization across different protein complexes. Beyond correlation metrics, Pythia-PPI also demonstrated superior prediction accuracy with the lowest RMSE of 1.08 and MAE of 0.738 among all methods, and achieved the highest discriminative power with an AUROC of 0.779. These results validate that our integrated approach combining transfer learning, multi-task learning, and self-distillation effectively captures the underlying patterns in protein-protein interactions.

**Table 1.**
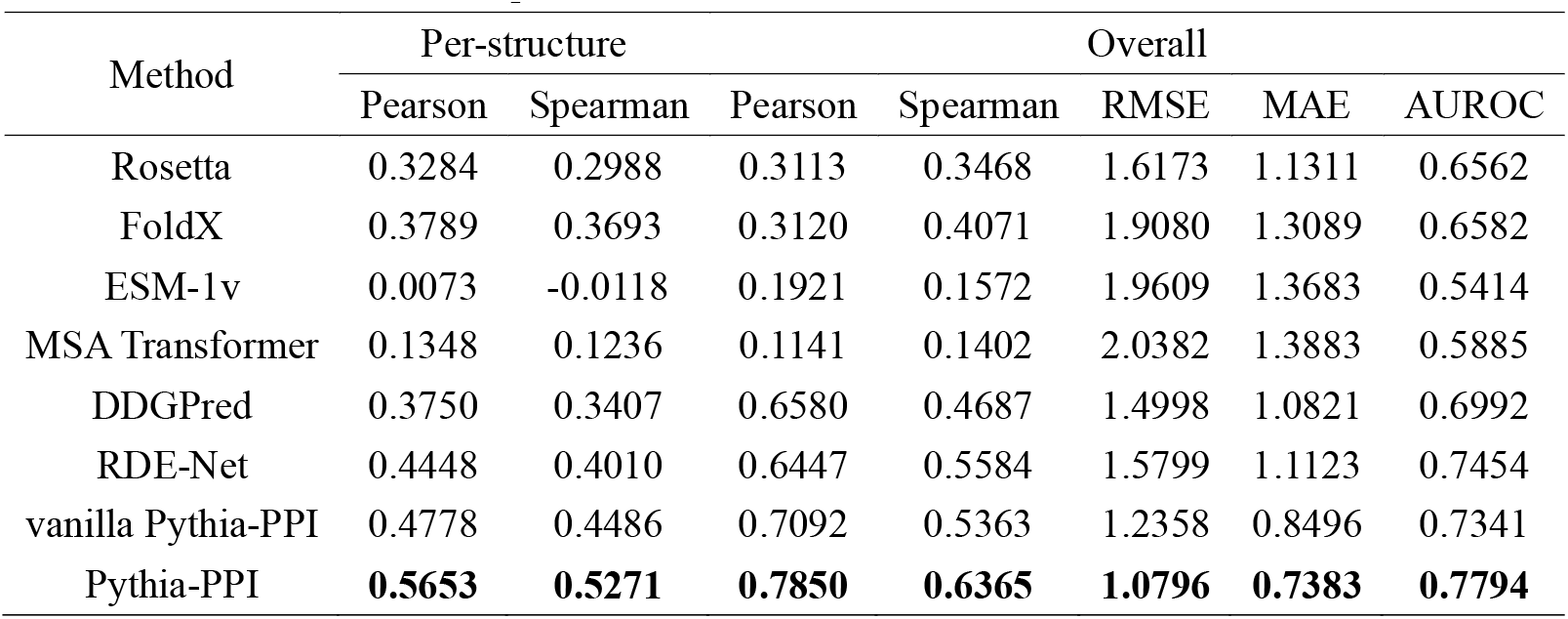
Comparison with various methods for the SKEMPI dataset.

Through ablation studies examining the model development pipeline, we observed that self-distillation contributed substantially to model performance (Figure 4E). Compared to the vanilla Pythia-PPI model without self-distillation, the final model achieved an absolute improvement of 0.10 in Spearman correlation. More notably, when compared to Pythia’s zero-shot predictions, the final Pythia-PPI model demonstrated a remarkable improvement with an absolute increase of 0.41 in Spearman correlation, highlighting the effectiveness of our multi-task and data augmentation strategy.

### Accurate prediction of viral protein fitness and antibody-antigen affinity from complex structure

Examination of RMSE in per-structure predictions revealed that the predictions predominantly maintained an RMSE below 2.0 kcal/mol regardless of the size of the complex structures (Figure 5A). The final model demonstrated strong anti-symmetry between direct and reversed mutation predictions, achieving a Pearson correlation of -0.9224 and Spearman correlation of -0.8926, indicating robust generalization despite extensive self-distillation training (Figure 5B). Structure-wise comparative analysis demonstrated that Pythia-PPI systematically outperformed Pythia across diverse protein complexes, maintaining superior or equivalent Pearson correlations in 97% of all evaluated structures (Figure 5C).

**Figure 5.**
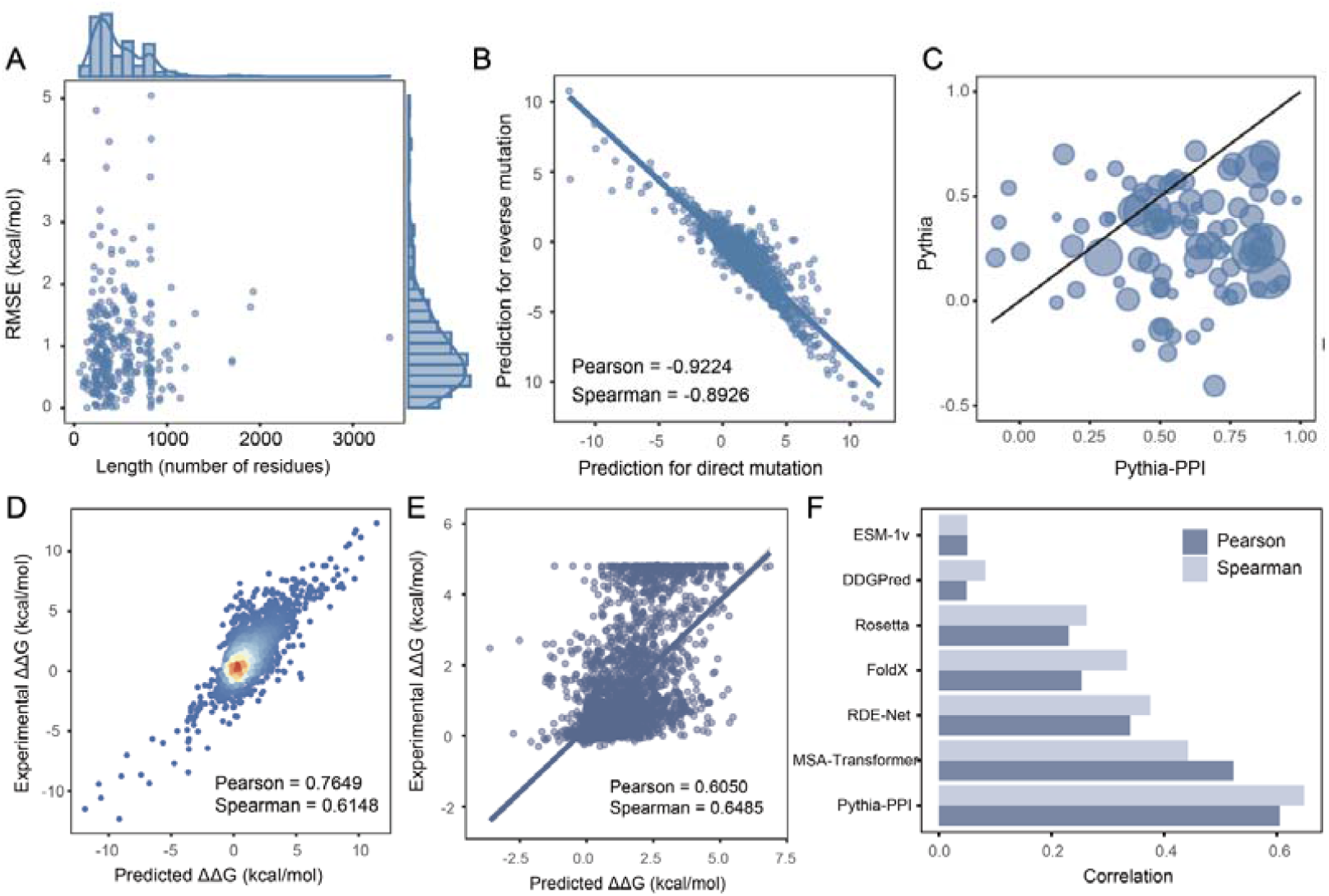
Performance analysis of Pythia-PPI on various datasets. (A) Scatter plot correlating per-structure length with the corresponding RMSE on the SKEMPI dataset. (B) Comparison of predictions for direct and reverse mutations. (C) Comparison of per-structure Pearson values of Pythia and Pythia-PPI. (D) Density scatter plot of Pythia-PPI predictions versus experimental values on the antibody-antigen subset of SKEMPI dataset. (E) Scatter plot of experimental and predicted ΔΔG on the R3669 dataset. (F) Bar plot of Pearson and Spearman correlation on R3669 of tested methods.

We further evaluated Pythia-PPI’s performance on antigen-antibody interactions, a crucial subset of PPIs involved in immune recognition and disease diagnosis. Analysis spanning 588 mutations across 27 antigen-antibody complexes from the SKEMPI dataset demonstrated strong predictive capability, with Pythia-PPI achieving a Pearson correlation of 0.6708 and Spearman correlation of 0.5263 (Figure 5D). We also tested Pythia-PPI on the deep mutation scanning dataset of the receptor binding domain of SARS-CoV-2 and human ACE2 receptor [31]. Using the reference structure from PDB entry 6M0J, we excluded terminal residues to create the R3669 dataset. Pythia-PPI demonstrated superior performance on this independent test set, achieving Pearson and Spearman correlations of 0.6051 and 0.6486, respectively (Figure 5E). These values markedly surpass the performance of previous state-of-the-art approaches (Table 2). By utilizing evolutionary information, MSA Transformer achieved the highest Pearson and Spearman correlations of 0.3654 and 0.4423. In the absence of evolutionary information, the structure-based deep learning model RDE-Net achieved Pearson and Spearman correlations 0.3395 and 0.3754 (Figure 5F). These results underscore the potential of Pythia-PPI in antibody optimization and viral protein fitness prediction.

**Table 2.** Comparison with other methods on the R3669 dataset.

| Method | Pearson | Spearman | RMSE | MAE | AUROC |
| --- | --- | --- | --- | --- | --- |
| Rosetta | 0.2305 | 0.2624 | 3.9869 | 3.1251 | 0.5970 |
| FoldX | 0.2533 | 0.3336 | 4.0439 | 2.2434 | 0.6293 |
| ESM-1v | 0.0505 | 0.0526 | 4.8661 | 4.5936 | 0.5230 |
| MSA Transformer | 0.3654 | 0.4423 | 7.8735 | 7.3045 | 0.7528 |
| DDGPred | 0.0492 | 0.0826 | 1.7700 | 1.0966 | 0.4932 |
| RDE-Net | 0.3395 | 0.3754 | 1.5700 | 0.9280 | 0.6516 |
| Pythia-PPI | **0.6051** | **0.6486** | **1.1834** | **0.8260** | **0.7956** |

## Discussion

Amino acid substitutions can impact the binding strength of protein-protein complexes, with some mutations stabilizing PPIs, while others weaken or even disrupt PPIs. This can influence the structure of protein-protein complexes and further affect their functions, potentially leading to the occurrence of diseases. Therefore, exploring the mutational effects on PPIs and achieving accurate and generalized predictions of ΔΔG values is of significant importance for protein engineering, disease treatment, and drug design. Through the integration of multi-task learning and self-distillation, Pythia-PPI achieved significant improvements in prediction accuracy. On the SKEMPI dataset, Pythia-PPI demonstrated enhanced performance with an overall Pearson correlation of 0.785. Notably, in per-structure evaluations, Pythia-PPI reached a Pearson correlation of 0.567, compared to 0.445 of previous state-of-the-art method, suggesting improved generalization across diverse protein complexes. Within subset of antibody-antigen interactions, Pythia-PPI’s predictive capability remained strong, achieving a Pearson correlation of 0.6708 and a Spearman correlation of 0.5263 on 588 mutations across 27 antibody-antigen complexes. Additionally, in the R3669 dataset focused on interactions between SARS-CoV-2 spike protein mutants and ACE2, Pythia-PPI displayed a notable increase in accuracy, achieving Pearson and Spearman correlations of 0.6051 and 0.6486, respectively. These results underline Pythia-PPI’s utility for applications in antibody optimization and viral protein mutation fitness prediction.

The design of the Pythia model was rooted in physical principles to predict protein thermostability, and it also generalized well to the prediction of protein binding affinity. The success of Pythia-PPI further emphasized the importance of designing artificial neural networks based on physical interpretations. The performance gains of Pythia-PPI can be attributed to the benefits of multi-task learning and the self-distillation strategy. By incorporating a protein stability prediction task, we leveraged the shared structural features between stability and binding affinity, enabling Pythia-PPI to capture a more comprehensive representation of protein structures, which should have enabled the model to generalize across more structures. The self-distillation process further utilized an augmented dataset from the predictions of the vanilla Pythia-PPI. This expanded dataset provided a more exhaustive mutation landscape, helping the model achieve a broader and more balanced distribution of examples, leading to substantial increases in both accuracy and robustness.

However, the self-distillation process introduced a minor bias in some predictions: compared to zero-shot predictions, the correlation of predicted and experimental values was decreased on approximately 3% of all structures in the SKEMPI dataset. This decrease indicates that the model may introduce specific biases due to self-distillation. The inherent limitations of model generated data were broadly discussed and the best way to develop better deep learning models relies on high-quality real data eventually. For examples, the recent published megascale dataset covering almost a million of mutations across hundreds of proteins [33], largely improved the prediction accuracy of protein stability [34]. A similar dataset for PPIs would be an invaluable resource for the development of robust deep learning models.

Beside using Pythia-PPI to predict the mutation effects on experimentally solved protein complex structures, the rapid advancements in structural prediction accuracy for protein complexes present new opportunities. With high-quality structural models of the human [35] and pathogen interactomes [36] becoming accessible recently, Pythia-PPI’s capability for rapid, evolution-free inference makes it particularly suitable to predict the fitness landscape of those important protein-protein interactions and would provide more insights into protein function and mechanisms.

## Methods and materials

### Dataset curation

In this study, we utilized three experimental datasets: the SKEMPI 2.0 dataset, the FireProt dataset, deep mutagenesis data on the spike protein of severe acute respiratory syndrome coronavirus 2 (SARS-CoV-2) in complex with human angiotensin-converting enzyme 2 (ACE2). The SKEMPI 2.0 dataset, containing PPI binding affinity data, and the FireProt dataset, comprising protein stability data, were used as training sets for the model. The deep mutagenesis dataset served as a blind test set to evaluate the model’s ability to predict changes in binding affinity due to mutations, while the S669 dataset assessed the model’s capability to predict stability changes induced by mutations. Comprehensive data cleaning was conducted on the SKEMPI 2.0, FireProt, and deep mutagenesis datasets to ensure data reliability. The SKEMPI 2.0 dataset underwent careful scrutiny, focusing on collecting single-point mutation samples and excluding entries with incomplete, uncertain, or unreliable data. Calculations were performed to determine the average change in binding affinity for identical single-point mutations across the same protein complexes, resulting in the SKEMPI dataset, which includes 4,076 single-point mutations across 314 protein-protein complexes. Mutations not present in the reference PDB structure (6M0J) of the SARS-CoV-2 spike protein, specifically those in the first two N-terminal residues and the last five C-terminal residues, were selectively excluded, leading to the filtered R3669 dataset. Adherence to the cleaning principles established by Dieckhaus et al. produced the F3436 dataset, comprising 3,436 single-point mutations from 100 proteins. The data were then transformed and integrated, extracting features required for the model, such as the PDB ID, unique identifiers of the protein-protein complexes, wild-type and mutant amino acids, mutation positions, and experimentally measured ΔΔG values.

### Dataset splitting

To ensure a robust and comprehensive assessment of the model’s performance, a protein-level 5-fold cross-validation strategy was implemented. Specifically, the SKEMPI dataset was divided into five folds based on protein-protein complex structures. In each iteration, each fold was designated as the validation set while the remaining folds were combined to form the training set.

This process was repeated five times, each time with a different fold used as the validation set, thereby ensuring a thorough and accurate evaluation. The average performance metrics from these five iterations were then used as the final evaluation metric for the model. Additionally, the F3436 dataset was entirely utilized for model training to enhance the model’s performance. Within the SKEMPI dataset splitting, the first fold included 841 mutation samples involving 75 protein-protein complexes; the second fold contained 819 mutation samples involving 56 protein-protein complexes; the third fold comprised 980 mutation samples involving 53 protein-protein complexes; the fourth fold included 819 mutation samples involving 91 protein-protein complexes; and the fifth fold encompassed 617 mutation samples involving 39 protein-protein complexes.

### Evaluation metrics

We utilized five metrics to assess the overall performance of the model, which include the Pearson correlation (Pearson correlation coefficient, r), Spearman correlation (Spearman correlation coefficient, ρ), RMSE (Root Mean Square Error), MAE (Mean Absolute Error), and AUROC (Area Under the Receiver Operating Characteristic Curve).

The formulas for these metrics are as follows:

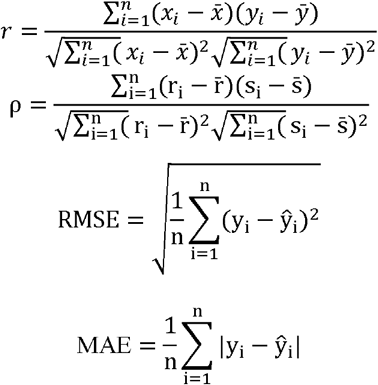

These metrics were employed to analyze the relationship between predicted and experimentally determined ΔΔG values. In practical applications, the correlation for specific protein-protein complexes often receives more attention. Consequently, for datasets containing multiple protein complexes, we grouped the mutation samples by structure following principles established by Luo et al. Groups containing fewer than ten mutation samples were discarded. We calculated the correlation for each group and then obtained two additional metrics through averaging: the per-structure Pearson and Spearman correlation coefficients.

### Training of the Pythia-PPI

Our basic program language is Python 3.10.13, on which Pytorch 2.1.2 with a default random seed 2024. Unless otherwise specified, Pythia-PPI is trained using the Adam optimizer with a learning rate set at 1e-4 and epochs set at 100. The training loss is computed using the absolute value error loss function. We employ a callback function (ReduceLROnPlateau) to update the learning rate, with a patience setting of 5 and a decay factor of 0.1. To prevent overfitting, early stopping is implemented with a patience setting of 10. All knowledge transfer experiments utilize the ‘pythia-p.pt’ model from Pythia, which the pretraining stage involved protein complexes. The batch size is set at 32, and all batches are sampled in each epoch. At the end of training, the model from the training fold with the highest diversity of protein-protein complexes is selected as the best model.

In the Pythia-PPI model, which addresses two distinct tasks, the loss functions are defined for each task respectively. The loss function for the PPI binding affinity prediction task is denoted as L_affinity_, while the loss function for the protein stability prediction task is denoted as L_stability_. The overall loss function for the model is expressed as:

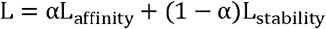

In obtaining the optimal model for PPI binding affinity, the loss weight α was set at 0.8. To stabilize the training process and enhance the model’s generalization ability on the protein stability dataset, the learning rate is set at 1e-7 and epochs at 1000. Training the Pythia-PPI model on the SKEMPI 2.0 dataset typically requires about half an hour on a single V100 GPU, supported by 10 dedicated CPUs.

### Data augmentation on SEKMPI dataset

To create the augmented dataset, we selected residues located on the protein-protein interface across all structures in the SKEMPI dataset. Residue pairs located on different chains and the distance of Ca within 7Å were identified as interface residues. The ΔΔG values of all possible mutations were predicted with the vanilla Pythia-PPI as augmented data and used to train the final

Pythia-PPI model following the same setting as the training of the vanilla Pythia-PPI.

### Baseline Methods

We utilized six different tools to predict ΔΔG values on the R3669 dataset, using the default model weights and inference processes for all methods. Here is a brief overview of their characteristics:

**Rosetta** [10] is a physics-based method that evaluates the stability of protein structures using an energy function. It combines Monte Carlo sampling algorithms to explore structural space and fragment assembly techniques to accurately predict and design protein structures.

**FoldX** [11] is another physics-based method that predicts protein structure and stability by calculating the interaction energy between individual atoms within the protein and between different amino acids.

**ESM-1v** [28] is a machine learning-based language model capable of zero-shot prediction, which allows it to predict the impact of protein mutations on function without specific training data.

**MSA Transformer** [29] is a machine learning-based language model that uses the self-attention mechanisms of Transformers to capture dependencies between sequences. It enhances protein sequence representation by modeling multiple sequence alignment data.

**DDGPred** [**Error! Reference source not found**.] is a supervised machine learning method that uses an attention-based geometric neural network structure to learn the geometric effects of protein-protein interactions from three-dimensional protein-protein complex structures.

**RDE-Net** [30] is also a machine learning method that utilizes unsupervised learning with Rotamer Density Estimation (RDE) by simulating the conformations of side-chain atoms to capture atomic interactions and learn rotameric density representations.

These tools provide a comprehensive suite of methods for assessing and predicting the effects of mutations in PPI, leveraging both traditional physics-based techniques and cutting-edge machine learning-based models.

## Notes

### Competing Interest Statement

The authors have declared no competing interest.

