## Supplementary material for "Reliable prediction of protein-protein binding affinity changes upon mutations with Pythia-PPI": SI

### Supplementary Figures

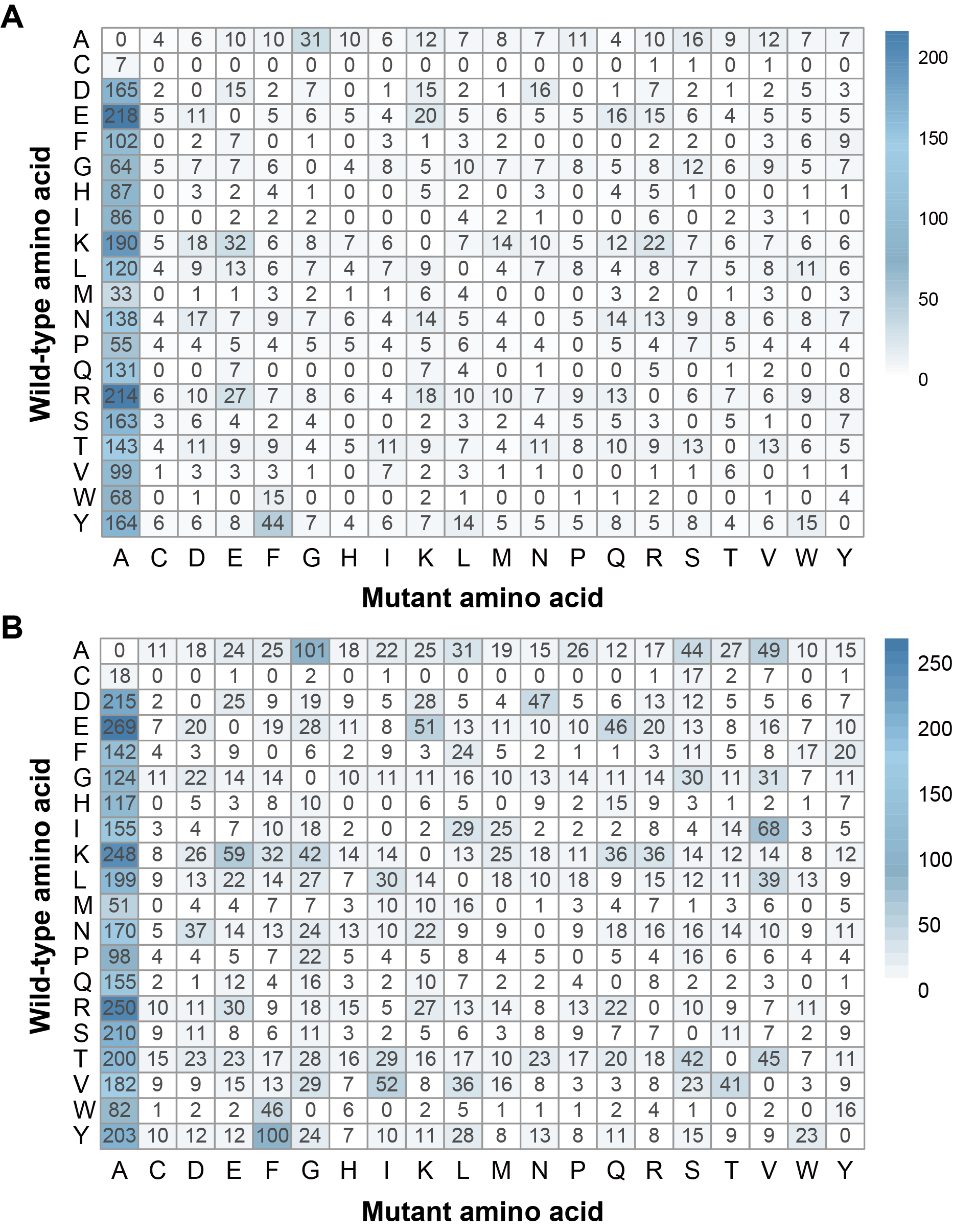

**Supplementary Figure 1.** (A) Wild-type and mutant amino acids for single-point mutations based on the SKEMPI dataset. (B) Wild-type and mutant amino acids for single-point mutations based on both the SKEMPI and the FireProt datasets.

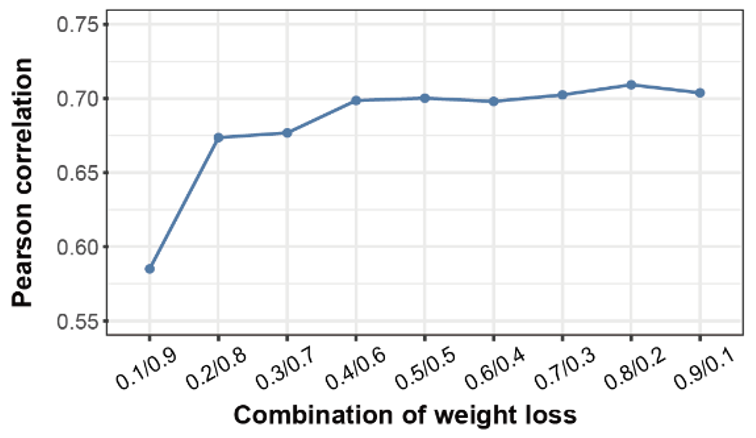

**Supplementary Figure 2.** Pearson correlation on the SKEMPI 5-fold cross validation of different loss weight combination.

### Supplementary Tables

**Supplementary Table 1.** Performance comparison of pre-trained and fine-tuned models for the S4076 dataset

| Method | Per-structure | | Overall | | | | |
| --- | --- | --- | --- | --- | --- | --- | --- |
|  | Pearson | Spearman | Pearson | Spearman | RMSE | MAE | AUROC |
| MIF-Δlogit | 0.2580 | 0.2364 | 0.2726 | 0.2658 | 7.7037 | 6.7320 | 0.6156 |
| Pythia-Δlogit | 0.2879 | 0.2579 | 0.3782 | 0.3036 | 5.9397 | 4.3640 | 0.6249 |
| MIF_FT | 0.4351 | 0.4059 | 0.6570 | 0.5045 | 1.3472 | 0.9067 | 0.7198 |
| Pythia_FT | 0.4462 | 0.4119 | 0.6939 | 0.5319 | 1.2687 | 0.8577 | 0.7286 |
| Pythia-PPI | **0.4778** | **0.4486** | **0.7092** | **0.5365** | **1.2358** | **0.8496** | **0.7341** |

### Supplementary Notes

#### 1. Generation of mutant protein-protein complex structures

To investigate the mutant structures of the R3669 dataset (PDB ID: 6M0J) and the S4076 dataset, we employed the comparative modeling software MODELLER [1]. Utilizing the wild-type structure as a template and adhering to the default parameters of the software, we constructed models of the mutant protein-protein complexes. During this process, MODELLER reconstructed the side chains of the mutated residues and made minor adjustments to the backbone and side chains of the entire complex. These modifications helped eliminate spatial conflicts and optimized the atomic interactions with the new mutant residues.

#### 2. Details for MIF_FT

The MIF initially encodes amino acid sequences as node features in a graph. Subsequently, it employs a Graph Convolutional Network (GCN) to perform convolution operations on these node features, aiming to capture relational information among the nodes. Following this, MIF utilizes known protein structures as input, converting structural information into graph edge features. The GCN is then reapplied to these edge features to extract interaction information between residues. The MIF model is pretrained to obtain feature representations, and a multilayer perceptron is integrated to adapt these feature inputs. It is important to note that the amino acid sequence generation code embedded within MIF is solely designed for monomeric proteins. Modifications are required to adapt this code for generating sequences of protein complexes.

#### 3. Details for Pythia_FT

Pythia is an autonomous graph neural network that integrates attention mechanisms with message passing neural networks (MPNN), focusing on identifying critical substructures during the learning process. These substructures are essential for understanding interaction properties. In Pythia, each amino acid is treated as a node, connected based on the Euclidean distance to the nearest 32 amino acids from the C-α atom. The model utilizes one-hot encoding to represent each amino acid type and employs sine and cosine functions to express main-chain dihedral angles (φ, ψ, and ω) as node input features. For edge features, Pythia considers the distances between five main-chain atoms (C-α, C, N, O, and C-β), along with their sequence positions and chain information. Through pretraining the Pythia model, we obtain feature representations and incorporate a multilayer perceptron to adapt these feature inputs. It is important to note that extracting corresponding wild-type and mutant features based on amino acid types and mutation positions is essential for feature extraction.

#### 4. Comprehensive performance comparison of pre-trained and fine-tuned models

We employed seven evaluation metrics to comprehensively compare the performance of MIF-Δlogit, Pythia-Δlogit, MIF_FT, Pythia_FT, and Pythia-PPI. The observations indicate that, compared to the pre-trained models, the fine-tuned models achieved an improvement of over 80% in overall Pearson correlation and more than 50% in per-structure correlation. Furthermore, both fine-tuned Pythia_FT compared to MIF_FT and Pythia-Δlogit relative to MIF-Δlogit outperformed across all evaluation metrics, demonstrating that representations derived from Pythia are indeed superior to those from MIF. Additionally, Pythia-PPI, built upon the foundation of Pythia_FT, further enhances performance comprehensively, particularly notable in terms of per-structure correlation.

#### 5. Detailed performance comparison with other methods on the S669 Dataset

Pythia-PPI consistently demonstrates superior performance across both the direct mutation set S669-DIR and the reverse mutation set S669-REV [2]. For S669-DIR, Pythia-PPI achieves a Pearson correlation of 0.4782 and a Spearman correlation of 0.4975, indicating strong agreement between predicted and experimental ΔΔG values. Its RMSE and MAE values are 1.5200 and 1.1000, respectively, reflecting high predictive accuracy and low error. In S669-REV, Pythia-PPI maintains robust performance with a Pearson correlation of 0.4743 and a Spearman correlation of 0.4679. The RMSE and MAE values for these reverse mutations are 1.8201 and 1.3885, respectively, showing slightly higher error but still strong predictive capability. Compared to other methods, such as MAESTRO, which performs well in S669-DIR but drops significantly in S669-REV, Pythia-PPI demonstrates consistent reliability. The consistent performance of Pythia-PPI across various evaluation metrics underscores its effectiveness in capturing the impacts of point mutations on protein-protein interactions.

**Supplementary Table 2.** Performance comparison of fourteen methods for the S669 dataset

| Method | S669-DIR | | | | S669-REV | | | | |
| --- | --- | --- | --- | --- | --- | --- | --- | --- | --- |
|  | Pearson | Spearman | RMSE | MAE | | Pearson | Spearman | RMSE | MAE |
| vanilla Pythia-PPI | 0.4782 | **0.4975** | 1.5200 | 1.1000 | | **0.4743** | **0.4679** | 1.8201 | 1.3885 |
| MAESTRO [3] | **0.4967** | 0.4634 | **1.4444** | 1.0619 | | 0.1977 | 0.1931 | 2.0972 | 1.6464 |
| ACDC-NN [4] | 0.4597 | 0.4540 | 1.4884 | **1.0454** | | 0.4505 | 0.4422 | 1.5048 | **1.0560** |
| DDGun3D [5] | 0.4324 | 0.4271 | 1.5971 | 1.1091 | | 0.4105 | 0.4106 | 1.6184 | 1.1438 |
| INPS3D [6] | 0.4303 | 0.4390 | 1.5028 | 1.0683 | | 0.3279 | 0.3585 | 1.7647 | 1.3083 |
| Pythia | 0.4292 | 0.4637 | 8.3886 | 6.1554 | | 0.4331 | 0.4453 | 6.3759 | 4.7321 |
| Dynamut [7] | 0.4148 | 0.3737 | 1.5955 | 1.1851 | | 0.3444 | 0.3649 | 1.6948 | 1.2378 |
| PoPMuSiC [8] | 0.4146 | 0.4149 | 1.5138 | 1.0879 | | 0.2395 | 0.2189 | 2.0856 | 1.6403 |
| DUET [9] | 0.4133 | 0.4160 | 1.5225 | 1.0962 | | 0.2283 | 0.2312 | 2.1412 | 1.6808 |
| SDM [10] | 0.4105 | 0.3920 | 1.6706 | 1.2644 | | 0.1348 | 0.1377 | 2.1562 | 1.6368 |
| PremPS [11] | 0.4050 | 0.4165 | 1.5128 | 1.0924 | | 0.4191 | 0.4210 | **1.4868** | 1.0541 |
| ThermoNet [12] | 0.3911 | 0.3733 | 1.6170 | 1.1742 | | 0.3785 | 0.3381 | 1.6573 | 1.2349 |
| mCSM [13] | 0.3593 | 0.3628 | 1.5438 | 1.1296 | | 0.2202 | 0.2086 | 2.3007 | 1.8598 |
| FoldX | 0.2141 | 0.2827 | 2.3179 | 1.5683 | | 0.2175 | 0.3390 | 2.4761 | 1.500 |
| I-Mutant3.0 [14] | 0.3589 | 0.3540 | 1.5366 | 1.1245 | | 0.1512 | 0.1537 | 2.3181 | 1.8678 |
